# Pronounced proliferation of non-beta cells in response to beta-cell mitogens in isolated human islets of Langerhans

**DOI:** 10.1101/2020.10.28.359158

**Authors:** Hasna Maachi, Julien Ghislain, Caroline Tremblay, Vincent Poitout

## Abstract

The potential to treat diabetes by increasing beta-cell mass is driving a major effort to identify beta-cell mitogens. Demonstration of mitogen activity in human beta cells is frequently performed in ex vivo assays. However, reported disparities in the efficacy of beta-cell mitogens led us to investigate the sources of this variability. We studied 35 male (23) and female (12) human islet batches covering a range of donor ages and BMI. Islets were kept intact or dispersed into single cells and cultured in the presence of harmine, glucose, or heparin-binding epidermal growth factor-like growth factor (HB-EGF), and subsequently analyzed by immunohistochemistry or flow cytometry. Proliferating cells were identified by double labelling with EdU and Ki67 and glucagon, c-peptide or Nkx6.1, and cytokeratin-19 to respectively label alpha, beta, and ductal cells. Harmine and HB-EGF stimulated human beta-cell proliferation, but the effect of glucose was dependent on the assay and the donor. Harmine potently stimulated alpha-cell proliferation and both harmine and HB-EGF increased proliferation of insulin- and glucagon-negative cells, including cytokeratin 19-positive cells. Given the abundance of non-beta cells in human islet preparations, our results suggest that assessment of beta-cell mitogens requires complementary approaches and rigorous identification of cell identity using multiple markers.

## INTRODUCTION

Type 2 diabetes is characterized by insufficient functional β-cell mass ^1^. In non-diabetic obese individuals and during pregnancy, accretion of insulin secretory capacity per β cell combined with β-cell mass expansion maintains glucose homeostasis by balancing levels of circulating insulin and insulin sensitivity. Deciphering the mechanisms controlling β-cell proliferation has thus become a major research goal in the hope to identify therapeutic targets to expand β-cell mass and prevent or delay the onset of diabetes.

Diverse factors including insulin ^2^, GLP-1 ^3^, serotonin ^4^, epidermal growth factors ^5^, osteoprotegerin ^6^, serpinB1 ^7^ and nutrients such as glucose and fatty acids ^8–12^ promote replication of rodent β cells. Unfortunately, our knowledge of the factors controlling human β-cell proliferation is much more limited ^13–15^. Although human β cells respond to glucose ^16–18^, fatty acids ^12^, serpinB1 ^7^, dual-specificity tyrosine-regulated kinase 1A (DYRK1A) inhibitors including harmine ^19–21^, and heparin-binding epidermal growth factor-like growth factor (HB-EGF) ^22^, proliferation rates are very low compared to rodents and published data on the efficacy of human β-cell mitogens are often highly variable ^23^.

In this study we assessed the mitogenic properties of harmine, glucose and HB-EGF in both intact and dispersed human islets comparing immunochemistry to flow cytometry to differentiate proliferating β and non-β cell types, including glucagon^+^ and cytokeratin-19 (CK19)^+^ cells.

## RESULTS

### Proliferative effects of harmine, glucose, and HB-EGF in dispersed human islets as assessed by immunocytochemistry

As the majority of previous studies examined the effects of mitogens on human β-cell proliferation in dispersed islet cultures ^17–19,21,24–27^, we began by attempting to reproduce these findings. Following recovery, human islets were dispersed, plated and exposed to harmine (10 μM), high glucose (16.7 mM) or HB-EGF (100 ng/ml) for 3 days in the presence of the proliferation marker 5-ethynyl-2’-deoxyuridine (EdU). At the end of the treatment, cells were stained for EdU and the nuclear marker of mature β cells Nkx6.1 ^23^ (Fig. 1). Exposure to harmine in the presence of 5.5 mM glucose lead to a significant increase in β-cell proliferation compared to vehicle in 5/6 donors (Fig. 1a-c) (fold-change 13.0 ± 4.0, b & c: P = 0.034). In contrast, 16.7 mM glucose was a poor inducer of β-cell proliferation compared to 2.8 mM glucose with only 2/6 donors responding (Fig. 1d, e & g) (fold-change 1.2 ± 0.4, e & g: NS). Exposure to HB-EGF in the presence of 2.8 mM glucose increased β-cell proliferation compared to vehicle in 5/6 donors (Fig. 1d, f & g) (fold-change 2.1 ± 0.5, f: P = 0.011, g: NS). Interestingly, we frequently observed Nkx6.1^-^/EdU^+^ cells in the presence of harmine (Fig. 1a) and HB-EGF (Fig. 1d), suggesting that non-β cells also respond to these compounds.

**Figure 1.**
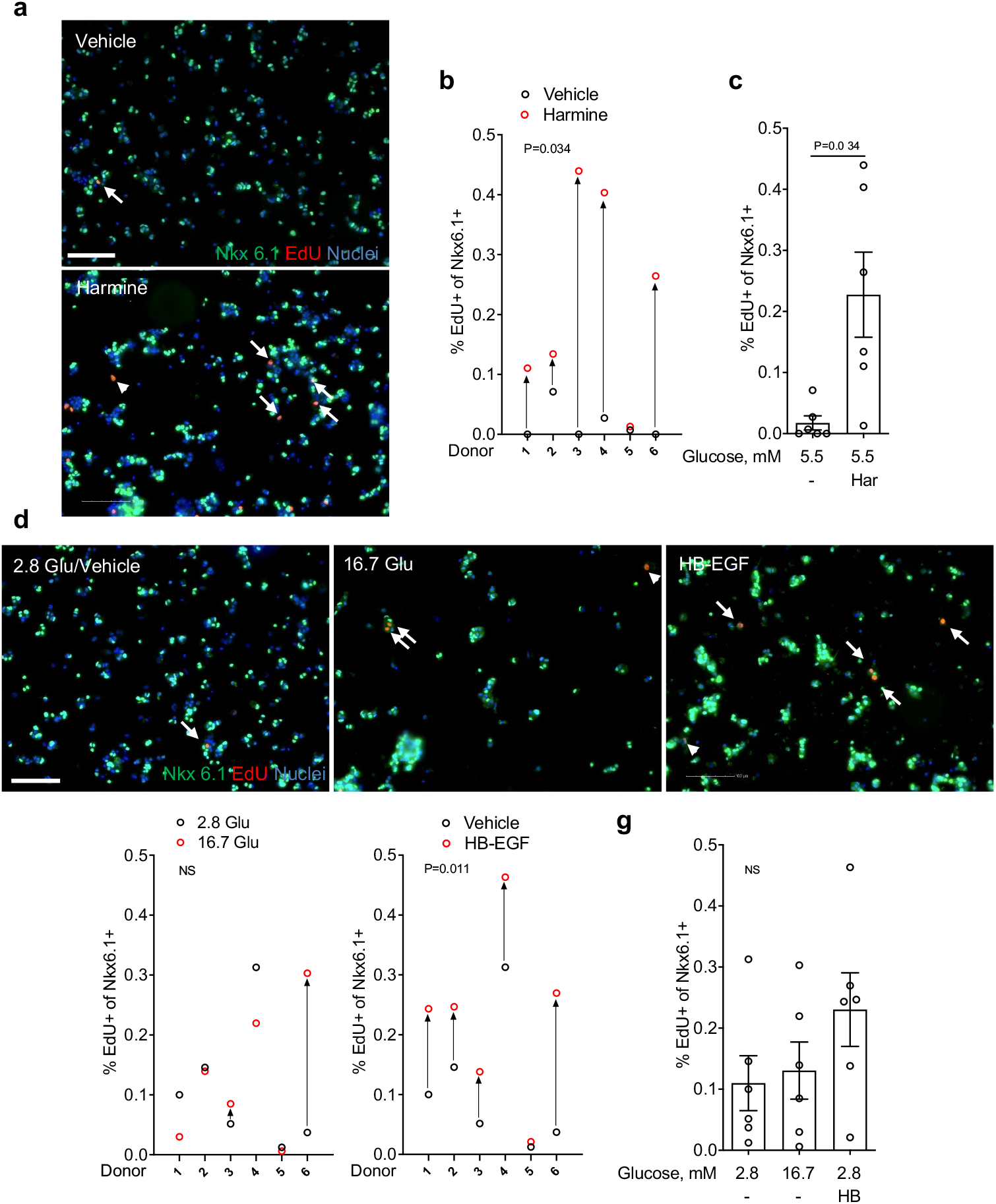
Analysis of β-cell proliferation in dispersed human islets by immunocytochemistry for Nkx6.1. (a-g) Dispersed human islets were exposed to 5.5 mM glucose (Vehicle; a-c), 2.8 mM glucose (Vehicle or 2.8 Glu; d-g), harmine (10 μM) (Har; a-c), 16.7 mM glucose (16.7 Glu; d, e & g) and HB-EGF (100 ng/ml) (HB; d, f & g) for 72 h. EdU (10 μM) was added throughout. Proliferation was assessed by EdU staining and Nkx6.1 to mark β cells. Representative images of Nkx6.1 (green), EdU (red) and nuclei (blue) staining are shown (a & d). Arrows and arrowheads highlight EdU^+^/Nkx6.1^+^ and EdU^+^/Nkx6.1^-^ cells, respectively. Scale bar, 100 μm. Images were acquired using an Operetta CLS with Harmony 4.5 software (https://www.perkinelmer.com/en-ca/product/operetta-cls-system-hh16000000). Between 1,000 and 1,500 Nkx6.1+ cells were counted for each sample. Proliferation is presented as the percentage of EdU^+^/Nkx6.1^+^ cells over the total Nkx6.1^+^ cells. Graphs showing cell proliferation of individual (b, e & f) and combined (c & g) donors were generated using GraphPad Prism 9 software (https://www.graphpad.com/scientific-software/prism/). Donor identifiers (Supplementary Table S1 online) are indicated. Significance was tested using paired t-tests (b, c, e & f) or one-way ANOVA (g). P<0.05 was considered significant. NS, not significant.

### Proliferative effects of harmine, glucose, and HB-EGF in intact human islets as assessed by immunohistochemistry

Islet architecture and cell-to-cell contact may contribute to the capacity of β cells to proliferate ^28^. Therefore, we exposed intact human islets to harmine, glucose or HB-EGF and assessed β-cell proliferation in cryosections by immunohistochemistry for Nkx6.1 and EdU (Fig. 2). As intact islets may be subject to increased cell death during extended culture, we first confirmed that EdU was not staining apoptotic cells. Double staining of islets from 1 donor exposed to harmine, glucose or HB-EGF for EdU and the apoptosis marker FragEL revealed no significant colocalization (Supplementary Fig. S1 online) suggesting that EdU^+^ cells are not apoptotic. Harmine increased proliferation of Nkx6.1^+^ cells in 5/8 donors (Fig. 2a, b & e) (fold-change 6.7 ± 2.2, b: P = 0.031, e: P = 0.031). In contrast to the results in dispersed islets, 7/9 donors responded to glucose (Fig. 2a, c & e) (fold-change 3.8 ± 1.0, c: P = 0.022, e: NS), and whereas 6/9 donors responded to HB-EGF the average effect was not significant (Fig. 2a, d & e) (foldchange 4.5 ± 2.1, d & e: NS). As observed in dispersed islets, many proliferating cells did not stain for Nkx6.1, which led us to quantify proliferation in this presumably non-β cell population. In basal conditions (2.8 mM glucose) the mean percentage of EdU^+^/Nkx6.1^-^ cells of the 9 donors was 1.3 ± 0.3 % (Fig. 2a & f-i). Harmine exposure led to a large increase in Nkx6.1^-^ cell proliferation in 8/8 donors (Fig. 2a, f & i) (fold-change 6.7 ± 2.0, f: P = 0.024, i: P = 0.0005). Glucose increased proliferation of Nkx6.1^-^ cells in 4/9 donors and this effect was not significant (Fig. 2a, g & i) (fold-change 1.2 ± 0.3, g & i: NS), whereas HB-EGF stimulated proliferation of Nkx6.1^-^ cells in 7/9 donors (Fig. 2a, h & i) (fold-change 1.6 ± 0.4, h: P = 0.01, i: NS).

**Figure 2.**
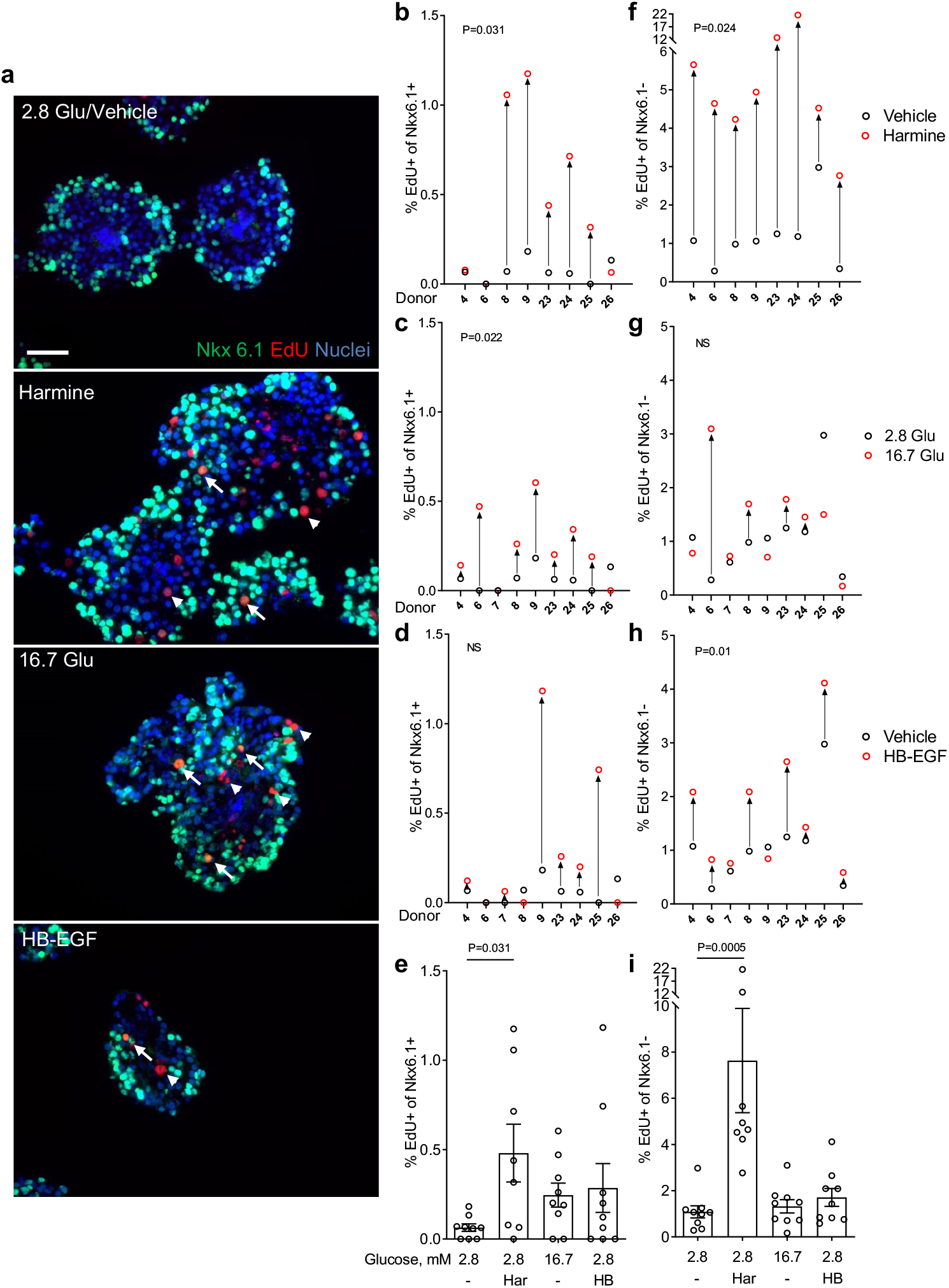
Analysis of β-cell proliferation in intact human islets by immunohistochemistry for Nkx6.1. (a-i) Intact human islets were exposed to 2.8 mM glucose (Vehicle or 2.8 Glu; a-i), harmine (10 μM) (Har; a & b, e, f & i), 16.7 mM glucose (16.7 Glu; a & c, e, g & i) and HB-EGF (100 ng/ml) (HB; a & d, e, h & i) for 72 h. EdU (10 μM) was added throughout. Proliferation was assessed by EdU staining and Nkx6.1 to mark β cells. Representative images of Nkx6.1 (green), EdU (red) and nuclei (blue) staining are shown (a). Arrows and arrowheads highlight EdU^+^/Nkx6.1^+^ and EdU^+^/Nkx6.1^-^ cells, respectively. Scale bar, 50 μm. Images were acquired using an Axio Imager with ZEN 2012 software (https://www.zeiss.com/microscopy/int/products/microscope-software/zen.html). A minimum of 1,500 Nkx6.1^+^ and Nkx6.1^-^ cells were counted for each sample. Proliferation is presented as the percentage of EdU^+^/Nkx6.1^+^ over the total Nkx6.1^+^ cells (b-e) or EdU^+^/Nkx6.1^-^ cells over the total Nkx6.1^-^ cells (f-i). Graphs showing cell proliferation of individual (b-d & f-h) and combined (e & i) donors were generated using GraphPad Prism 9 software (https://www.graphpad.com/scientific-software/prism/). Donor identifiers (Supplementary Table S1 online) are indicated. Significance was tested using paired t-tests (b-d & f-h) or one-way ANOVA (e & i). P<0.05 was considered significant. NS, not significant.

To confirm the results obtained with Nkx6.1 in intact islets, we used C-peptide (CPEP) as another β cell marker (Fig. 3). Similar to the results shown in Fig. 2, harmine, glucose and HB-EGF increased proliferation of CPEP^+^ cells in 7/7 donors (Fig. 3a, b & e) (fold-change 6.4 ± 1.7, b: P = 0.025, e: P = 0.0013), 5/6 donors (Fig. 3a, c & e) (fold-change 2.0 ± 0.5, c: P = 0.021, e: NS) and in 6/7 donors (Fig. 3a, d & e) (fold-change 2.6 ± 0.6, d: P = 0.014, e: NS), respectively. Surprisingly, the fraction of EdU^+^/CPEP^+^ cells in both basal and stimulated conditions was considerably larger compared to EdU^+^/Nkx6.1^+^ cells from several donors (compare Fig. 3b-e to Fig. 2b-e). To confirm this observation, we stained donor islets (N = 3) simultaneously for CPEP, Nkx6.1 and EdU (Supplementary Fig. S2 online). On average of all conditions, Nkx6.1^+^ cells represented 42 ± 2 % of the CPEP^+^ cells, whereas proliferating Nkx6.1^+^ cells represented only 13 ± 4 % of the proliferating CPEP^+^ cells. These data indicate that not all β-cells are detectable by Nkx6.1 staining and confirm our findings that more proliferating β cells are CPEP^+^ than Nkx6.1^+^. The proliferative response of CPEP^-^ cells to harmine (Fig. 3a, f & i), (6/7 donors; fold-change 9.7 ± 2.3, f: P = 0.012, i: P = 0.0007) glucose (Fig. 3a, g & i) (5/6 donors; foldchange 2.0 ± 0.4, g: P = 0.014, i: NS) and HB-EGF (Fig. 3a, h & i) (5/7 donors; fold-change 3.2 ± 1.4, h & i: NS) was similar to the response of Nkx6.1^-^ cells (Fig. 2).

**Figure 3.**
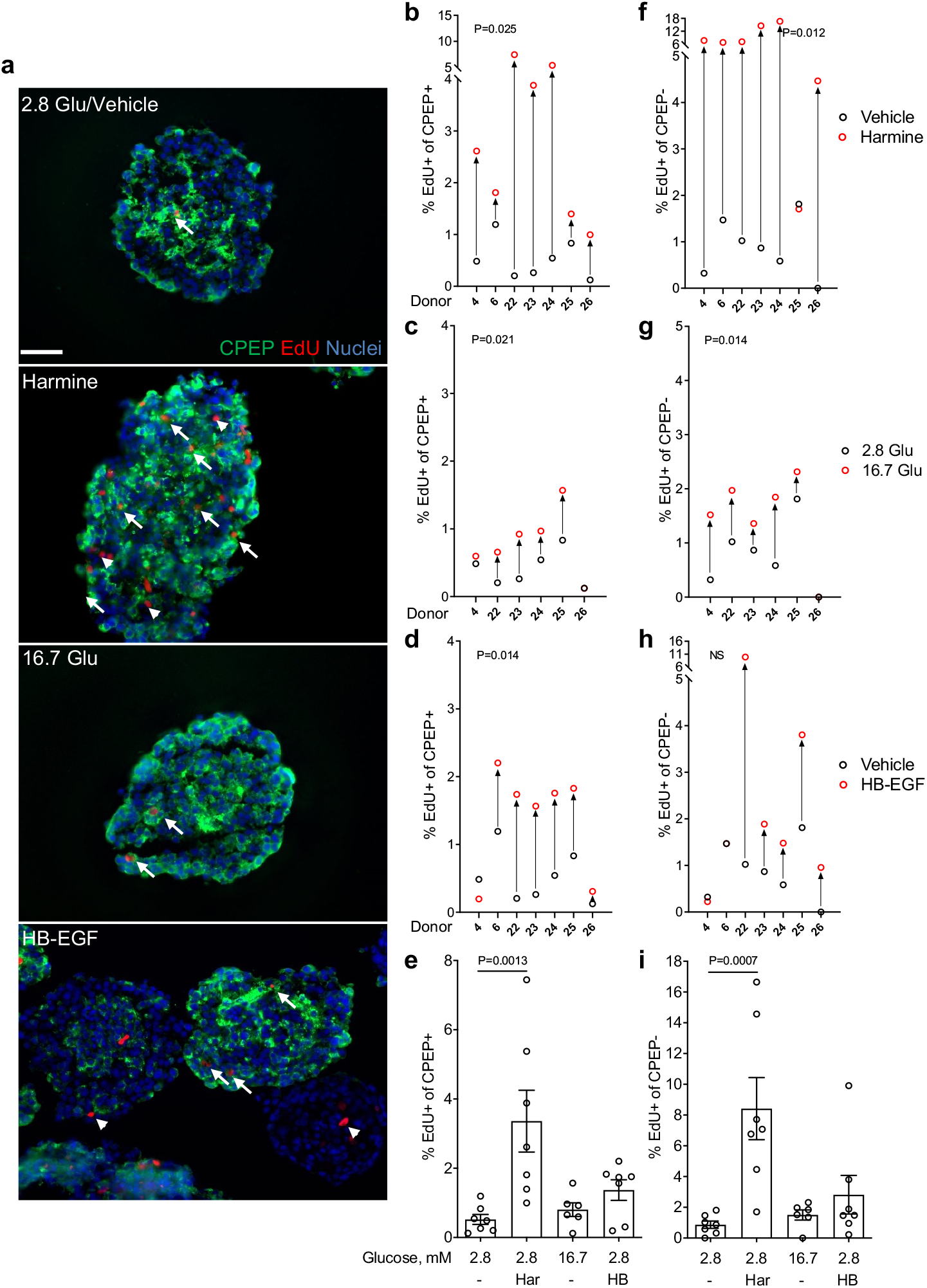
Analysis of β-cell proliferation in intact human islets by immunohistochemistry for C-peptide. (a-i) Intact human islets were exposed to 2.8 mM glucose (Vehicle or 2.8 Glu; a-i), harmine (10 μM) (Har; a & b, e, f & i), 16.7 mM glucose (16.7 Glu; a & c, e, g & i) and HB-EGF (100 ng/ml) (HB; a & d, e, h & i) for 72 h. EdU (10 μM) was added throughout. Proliferation was assessed by EdU staining and C-peptide (CPEP) to mark β cells. Representative images of CPEP (green), EdU (red) and nuclei (blue) staining are shown (a). Arrows and arrowheads highlight EdU^+^/CPEP^+^ and EdU^+^/CPEP^-^ cells, respectively. Scale bar, 50 μm. Images were acquired using an Axio Imager with ZEN 2012 software (https://www.zeiss.com/microscopy/int/products/microscope-software/zen.html). A minimum of 1,500 CPEP^+^ and CPEP^-^ cells were counted for each sample. Proliferation is presented as the percentage of EdU^+^/CPEP^+^ over the total CPEP^+^ cells (b-e) or EdU^+^/CPEP^-^ cells over the total CPEP^-^ cells (f-i). Graphs showing cell proliferation of individual (b-d & f-h) and combined (e & i) donors were generated using GraphPad Prism 9 software (https://www.graphpad.com/scientific-software/prism/). Donor identifiers (Supplementary Table S1 online) are indicated. Significance was tested using paired t-tests (b-d & f-h) or one-way ANOVA (e & i). P<0.05 was considered significant. NS, not significant.

To confirm the β-cell proliferative response to mitogens with a different proliferation marker we stained for the cell cycle antigen Ki67 and CPEP to mark β cells (Fig. 4). Similar to the results using EdU (Fig. 3), harmine, glucose and HB-EGF increased the percentage of Ki67^+^/CPEP^+^ cells in 5/5 donors (Fig. 4a, b & e) (fold-change 8.9 ± 2.2, b: P = 0.014, e: P = 0.002), in 4/5 donors (Fig. 4a, c & e) (fold-change 1.5 ± 0.4, c: P = 0.009, e: NS) and in 5/5 donors (Fig. 4a, d & e) (fold-change 4.2 ± 1.5, d & e: NS), respectively. Importantly, the percentage of Ki67^+^/CPEP^+^ detected under the different conditions was comparable to the percentage of EdU^+^/CPEP^+^ when donor islets were analyzed in parallel with both proliferation markers (compare Fig. 4 to Fig. 7 donors 30, 31 and 33). The percentage of Ki67^+^/CPEP^-^ cells increased in response to harmine in 5/5 donors (Fig. 4a, f & i), (fold-change 23.8 ± 8.8, f: P = NS, i: P = 0.0012), but only in 2/5 donors in response to glucose (Fig. 4a, g & i) (fold-change 1.4 ± 0.7, g & i: NS) and in 3/5 donors in response to HB-EGF (Fig. 4a, h & i) (fold-change 6.9 ± 4.2, h & i: NS) and was similar to the results obtained using EdU (Fig. 3). Overall, comparable results were obtained when using EdU or Ki67 as a proliferation marker of CPEP^+^ and CPEP^-^ in all conditions tested.

**Figure 4.**
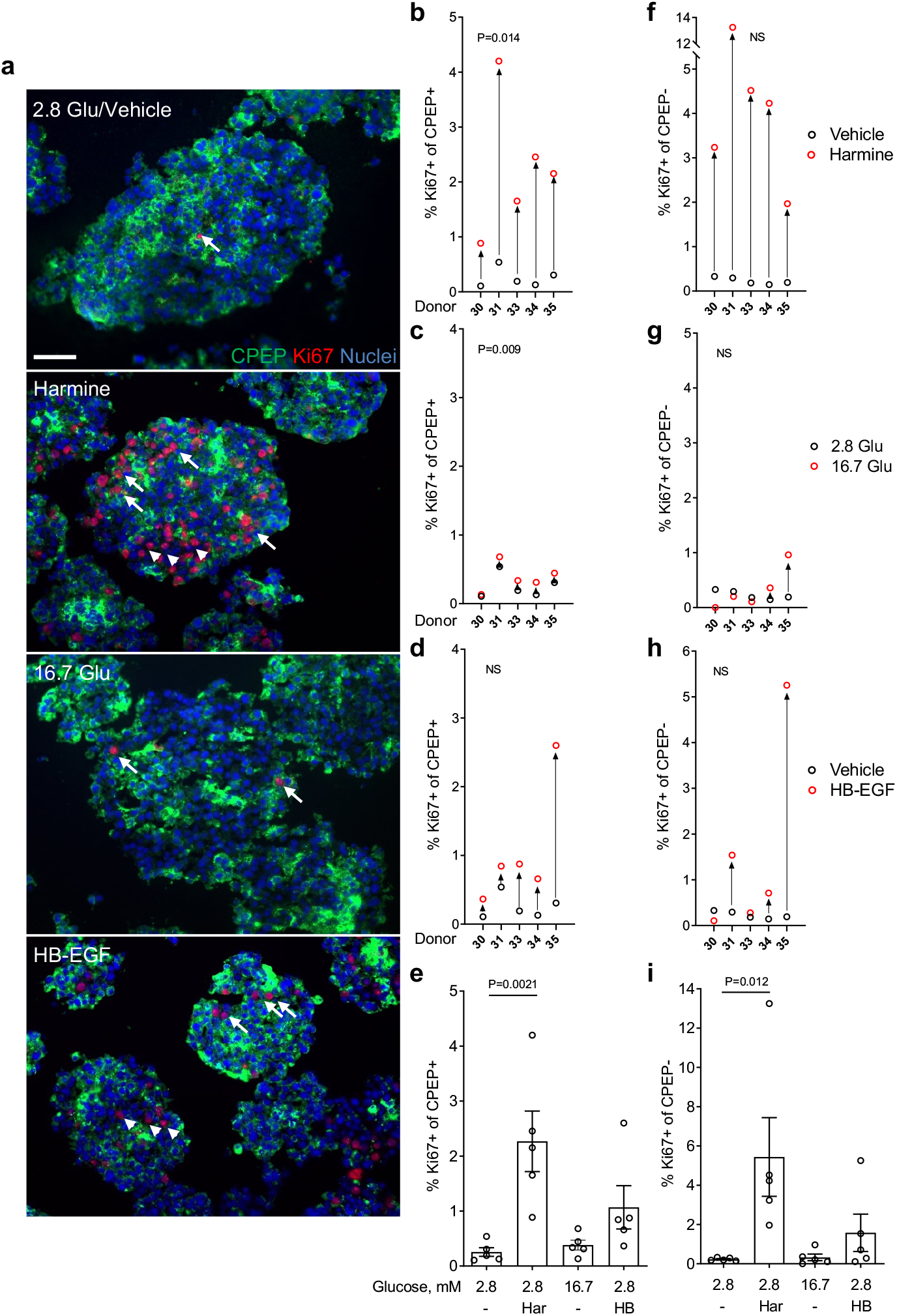
Analysis of β-cell proliferation in intact human islets by immunohistochemistry for Ki67. (a-i) Intact human islets were exposed to 2.8 mM glucose (Vehicle or 2.8 Glu; a-i), harmine (10 μM) (Har; a & b, e, f & i), 16.7 mM glucose (16.7 Glu; a & c, e, g & i) and HB-EGF (100 ng/ml) (HB; a & d, e, h & i) for 72 h. Proliferation was assessed by staining for Ki67 and C-peptide (CPEP) to mark β cells. Representative images of CPEP (green), Ki67 (red) and nuclei (blue) staining are shown (a). Arrows and arrowheads highlight Ki67^+^/CPEP^+^ and Ki67^+^/CPEP^-^ cells, respectively. Scale bar, 50 μm. Images were acquired using an Axio Imager with ZEN 2012 software (https://www.zeiss.com/microscopy/int/products/microscope-software/zen.html). A minimum of 1,500 CPEP^+^ and CPEP^-^ cells were counted for each sample. Proliferation is presented as the percentage of Ki67^+^/CPEP^+^ over the total CPEP^+^ cells (b-e) or Ki67^+^/CPEP^-^ cells over the total CPEP^-^ cells (f-i). Graphs showing cell proliferation of individual (b-d & f-h) and combined (e & i) donors were generated using GraphPad Prism 9 software (https://www.graphpad.com/scientific-software/prism/). Donor identifiers (Supplementary Table S1 online) are indicated. Significance was tested using paired t-tests (b-d & f-h) or one-way ANOVA (e & i). P<0.05 was considered significant. NS, not significant.

**Figure 5.**
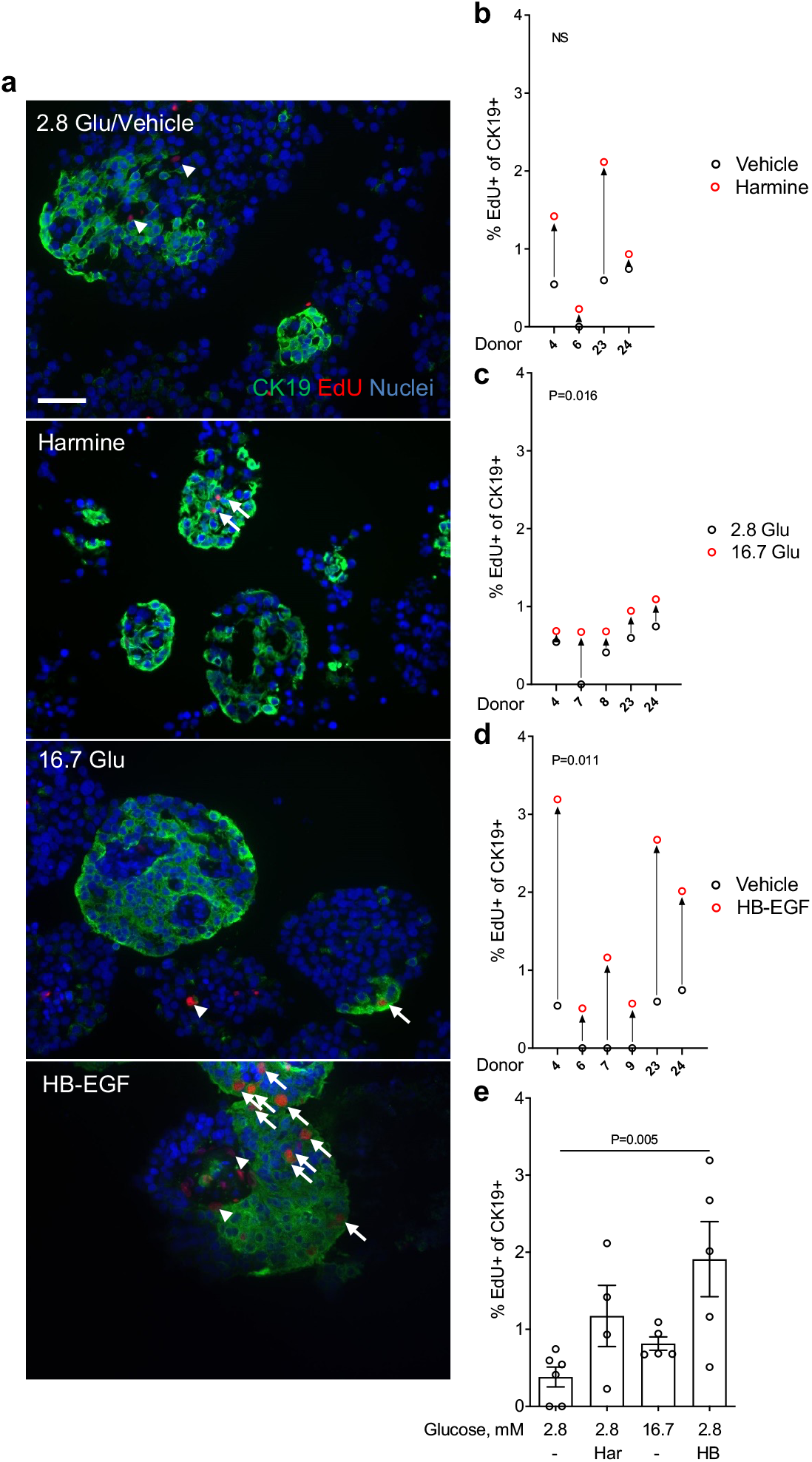
Analysis of cytokeratin 19-positive cell proliferation in intact human islets by immunohistochemistry. (a-e) Intact human islets were exposed to 2.8 mM glucose (Vehicle or 2.8 Glu; a-e), harmine (10 μM) (Har; a, b & e), 16.7 mM glucose (16.7 Glu; a, c & e) and HB-EGF (100 ng/ml) (HB; a, d & e) for 72 h. EdU (10 μM) was added throughout. Proliferation was assessed by EdU staining and cytokeratin 19 (CK19) to mark ductal cells. Representative images of CK19 (green), EdU (red) and nuclei (blue) staining are shown (a). Arrows and arrowheads highlight EdU^+^/CK19^+^ and EdU^+^/CK19^-^ cells, respectively. Scale bar, 50 μm. Images were acquired using an Axio Imager with ZEN 2012 software (https://www.zeiss.com/microscopy/int/products/microscope-software/zen.html). A minimum of 1,000 CK19^+^ cells were counted for each sample. Proliferation is presented as the percentage of EdU^+^/CK19^+^ over the total CK19^+^ cells (b-e). Graphs showing cell proliferation of individual (b-d) combined (e) donors were generated using GraphPad Prism 9 software (https://www.graphpad.com/scientific-software/prism/). Donor identifiers (Supplementary Table S1 online) are indicated. Significance was tested using paired t-tests (b-d) or one-way ANOVA (e). P<0.05 was considered significant. NS, not significant.

**Figure 6.**
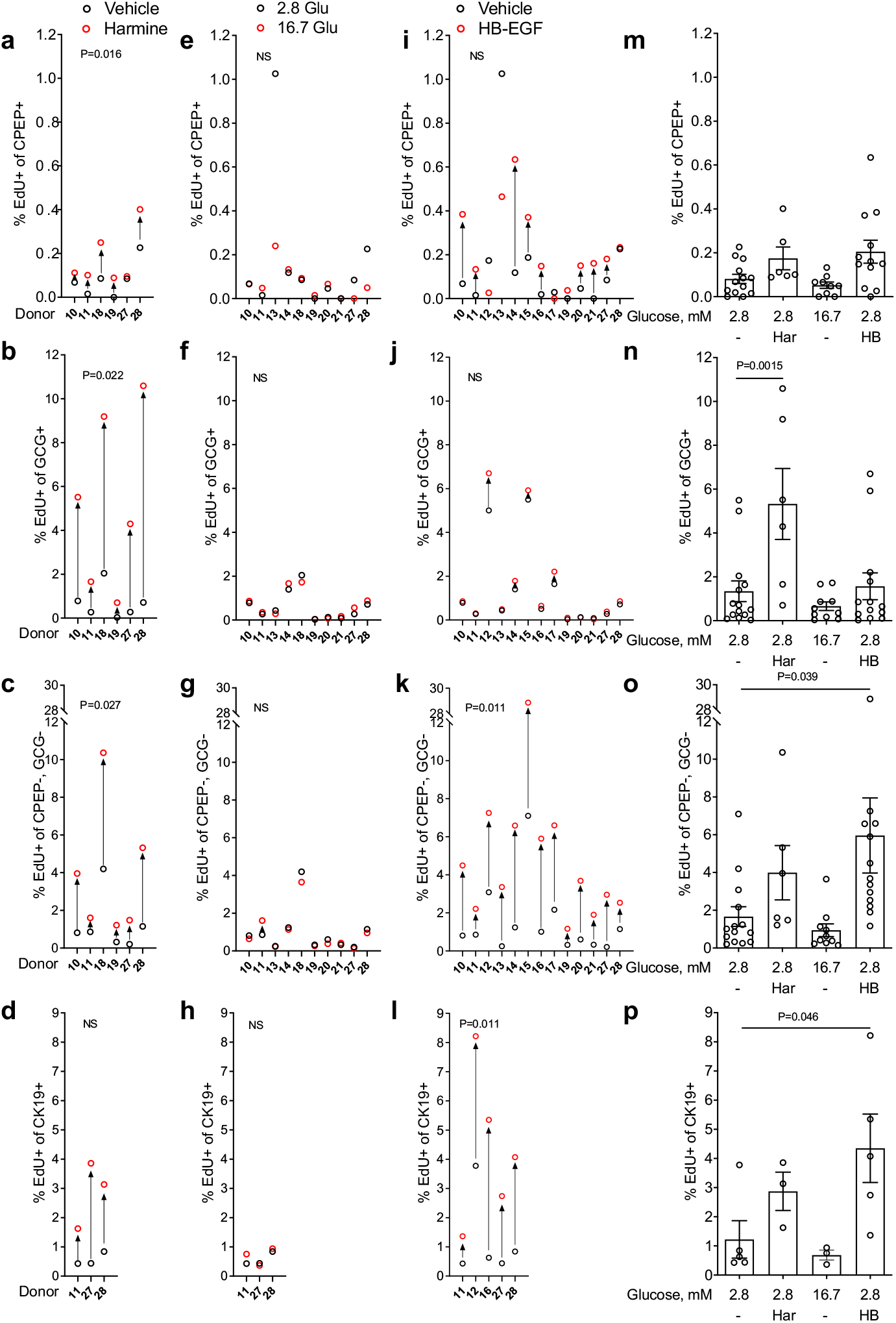
Analysis of cell proliferation in intact human islets by flow cytometry. (a-p) Intact human islets were exposed to 2.8 mM glucose (Vehicle or 2.8 Glu; a-p), harmine (10 μM) (Har; a-d & m-p), 16.7 mM glucose (16.7 Glu; e-h & m-p) and HB-EGF (100 ng/ml) (HB; i-l & m-p) for 72 h. EdU (10 μM) was added throughout. At the end of the treatment islets were dispersed into single cells and analyzed by flow cytometry for C-peptide (CPEP), glucagon (GCG), cytokeratin 19 (CK19) and EdU. A minimum of 2,000 CPEP^+^, GCG^+^, GCG^-^/CPEP^-^ and CK19^+^ cells were counted for each sample. Proliferation is presented as the percentage of EdU^+^ cells over the total cells for each of CPEP^+^ (a, e, i & m), GCG^+^ (b, f, j & n), CPEP^-^, GCG^-^ (c, g, k & o) and CK19^+^ (d, h, l & p) cell types. Graphs showing cell proliferation of individual (a-l) and combined (m-p) donors were generated using GraphPad Prism 9 software (https://www.graphpad.com/scientific-software/prism/). Donor identifiers (Supplementary Table S1 online) are indicated. Significance was tested using paired t-tests (a-l) and one-way ANOVA (m-p). P<0.05 was considered significant. NS, not significant.

**Figure 7.**
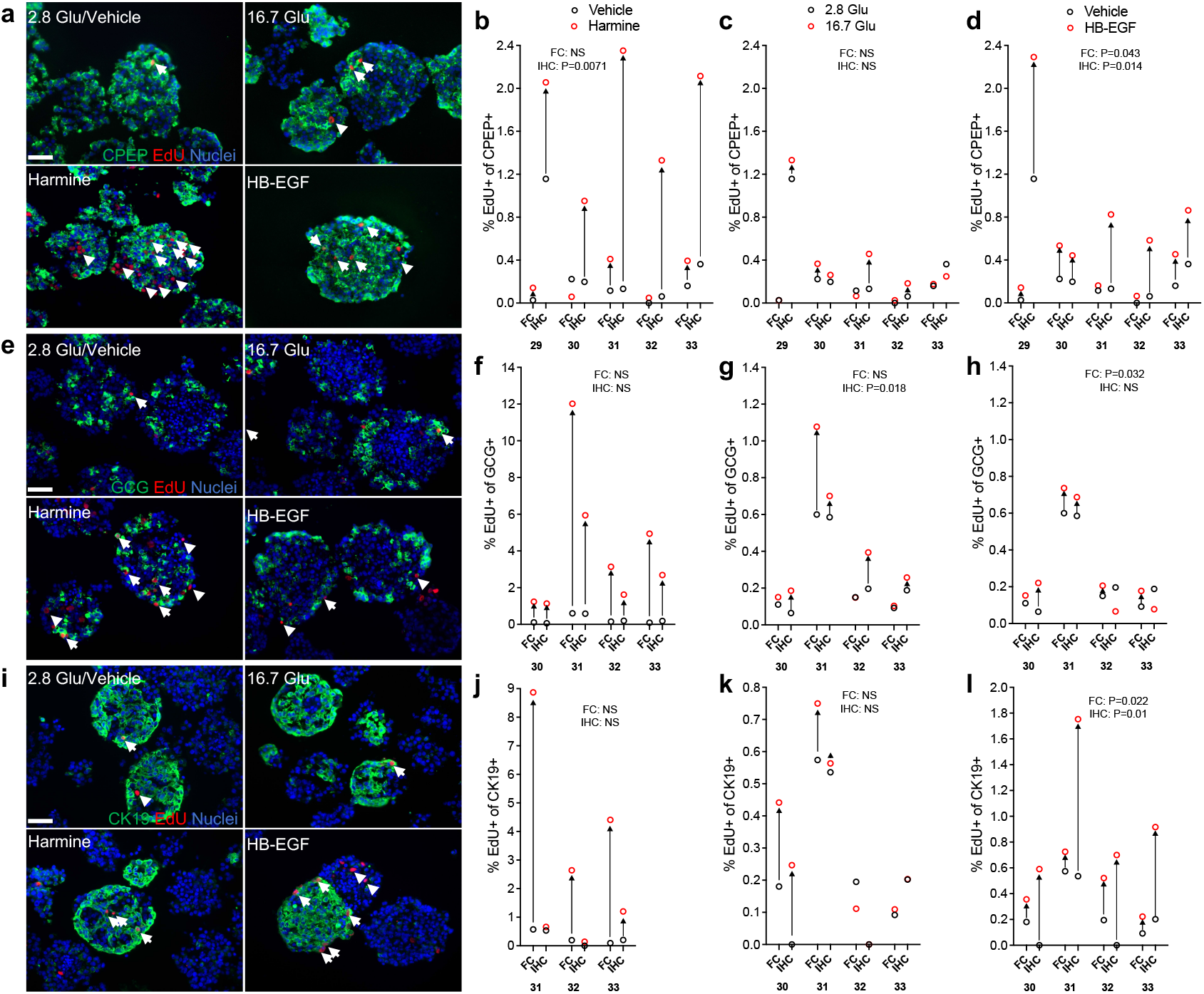
Comparison of immunohistochemistry and flow cytometry to assess human islet cell proliferation. (a-l) Intact human islets were exposed to 2.8 mM glucose (Vehicle or 2.8 Glu; a-l), harmine (10 μM) (a, b, e, f, i & j), 16.7 mM glucose (16.7 Glu; a, c, e, g, i & k) and HB-EGF (100 ng/ml) (a, d, e, h, i & l) for 72 h. EdU (10 μM) was added throughout. Proliferation was assessed in parallel for each donor by immunohistochemistry (IHC) and flow cytometry (FC) following staining for C-peptide (CPEP), glucagon (GCG), cytokeratin 19 (CK19) and EdU. (a, e & i) Representative images of CPEP (a, green), GCG (e, green), CK19 (i, green), EdU (red) and nuclei (blue) staining are shown. Arrows and arrowheads highlight respectively, EdU^+^/CPEP^+^ and EdU^+^CPEP^-^ (a), EdU^+^/GCG^+^ and EdU^+^/GCG^-^ (e) and EdU^+^/CK19^+^ and EdU^+^/CK19^-^ (i). Scale bar, 50 μm. Images were acquired using an Axio Imager with ZEN 2012 software (https://www.zeiss.com/microscopy/int/products/microscope-software/zen.html). A minimum of 1,000 and 2,000 CPEP^+^, GCG^+^ and CK19^+^ cells were counted for each sample in immunohistochemistry and flow cytometry, respectively. Proliferation is presented as the percentage of EdU^+^ cells over the total cells for each of CPEP^+^ (a-d), GCG^+^ (e-h) and CK19^+^ (il) cell types. Graphs showing cell proliferation of individual donors were generated using GraphPad Prism 9 software (https://www.graphpad.com/scientific-software/prism/). Donor identifiers (Supplementary Table S1 online) are indicated.

Human islet preparations often contain abundant non-endocrine cell types including duct and acinar cells, among others ^29^. The abundance of proliferating non-β cells (Fig. 2-4) led us to ask whether a proportion of these cells are positive for the ductal marker CK19 ^30^. Intact human islets were exposed to harmine, glucose or HB-EGF and β-cell proliferation assessed by immunohistochemistry for CK19 and EdU (Fig. 5). Numerous clusters of CK19^+^ cells were detected in all human islet preparations tested that were distinct from CPEP^+^ cells (Fig. 5a & Supplementary Fig. S3 online). Although harmine increased proliferation of CK19^+^ cells in 4/4 donors, the increase was not significant (Fig. 5a, b & e) (fold-change 2.5 ± 0.8, b & e: NS). In contrast glucose, and to a greater extent HB-EGF, increased CK19^+^ cell proliferation in 5/6 (Fig. 5a, c & e) (fold-change 1.8 ± 0.2, c: P = 0.016, e: NS) and 6/6 (Fig. 5a, d & e) (fold-change 5.4 ± 1.5, d: P = 0.011, e: P = 0.005) donors, respectively.

### Proliferative effects of harmine, glucose, and HB-EGF in intact human islets as assessed by flow cytometry

The reliability of immunohistochemical assessment of β-cell proliferation is inherently limited by the number of cells that can be manually counted and the operator’s visual appreciation of colabeling. We therefore exposed intact human islets to mitogens and assessed islet cell proliferation by flow cytometry using EdU and immunostaining for CPEP (β cells), glucagon (GCG; α cells) and CK19 (ductal cells) (Fig. 6 & Supplementary Fig. S4 online). On average, single cells represented 91 ± 3 % of total islet cells of which 88 ± 2 % were live cells (Supplementary Table S1 online). The mean proportion of β and α cells in the donor islet batches was 19 ± 3 % and 42 ± 5 %, respectively, whereas GCG^-^/CPEP^-^ cells represented the remaining 39 ± 5 % (Supplementary Table S1 online). Although stringent gating of CPEP^+^ cells may have led to a slight underestimation of the proportion of β-cells, these data are comparable to non-diabetic donors in the Human Pancreas Analysis Program (HPAP-RRID:SCR_016202) Database (https://hpap.pmacs.upenn.edu). Harmine induced β-cell proliferation in 5/6 donors (Fig. 6a & m) (fold-change 2.2 ± 0.6, a: P = 0.016, m: NS). Harmine was also a powerful mitogen for α cells in 6/6 donors (Fig. 6b & n) (fold-change 7.7 ± 2.4, b: P = 0.022, n: P = 0.0015) and increased proliferation of GCG^-^/CPEP^-^ cells in 6/6 donors (Fig. 6c & o) (foldchange 3.2 ± 1.1, c: P = 0.027, o: NS) and CK19^+^ cells in 3/3 donors (Fig. 6d & p) (fold-change 5.0 ± 1.1, d & p: NS). Glucose did not promote β-cell proliferation in any of the 10 donors tested (Fig. 6e & m) (fold-change 0.4 ± 0.1, e & m: NS). Furthermore, unlike harmine, glucose was ineffective as an α cell mitogen (Fig. 6f & n) (10 donors; fold-change 1.1 ± 0.3, f & n: NS), GCG^-^/CPEP^-^ cells (Fig. 6g & o) (10 donors; fold-change 0.9 ± 0.3, g & o: NS) and CK19^+^ cells (Fig. 6h & p) (3 donors; fold-change 1.2 ± 0.3, h & p: NS). HB-EGF increased β-cell proliferation in 9/13 donors (Fig. 6i & m) (fold-change 1.5 ± 0.3, i & m: NS), an effect that was significant (fold-change 2.5 ± 0.6, i: P = 0.027, m: P = 0.04) when donor 13, identified as an outlier based on Grubb’s test, was excluded from the analysis. HB-EGF was largely inactive on α cells (Fig. 6j & n) (13 donors; fold-change 1.2 ± 0.5, j & n: NS) but strongly stimulated GCG^-^ /CPEP^-^ cell proliferation in 13/13 donors (Fig. 6k & o) (fold-change 4.0 ± 1.3, k: P = 0.011, o: P = 0.039) and CK19^+^ cell proliferation in 5/5 donors (Fig. 6l & p) (fold-change 3.5 ± 1.0, l: P = 0.011, p: P = 0.046).

### Comparison of immunohistochemistry and flow cytometry to assess human islet cell proliferation in response to harmine, glucose, and HB-EGF

As variability in the proliferative response between batches of donor islets prevents the comparison of immunohistochemistry (IHC) and flow cytometry (FC), we performed these techniques in parallel on the same donor islets to investigate β-, α- and CK19^+^-cell proliferation. Intact human islets were exposed to harmine, glucose or HB-EGF and cell proliferation quantified by IHC in intact islet sections and by FC following staining for CPEP, GCG, CK19 and EdU (Fig. 7). Both techniques revealed a comparable percentage of proliferating CPEP^+^ cells at 2.8 mM glucose in 4/5 donors (Fig. 7a-d). Although a significant increase was detected in 5/5 donors by IHC in response to harmine (Fig. 7a & b) (fold-change 4.6 ± 0.7, P = 0.007), the response was much weaker when analyzed by FC (Fig. 7b) (fold-change 2.0 ± 0.8, NS). The response to 16.7 mM glucose was comparable in 4/5 donors (Fig. 7a & c) (FC: fold-change 1.2 ± 0.6, NS; IHC: fold-change 1.3 ± 0.6, NS), whereas the response to HB-EGF was similar in only 2/5 donors (Fig. 7a & d) (FC: fold-change 2.6 ± 0.9, P = 0.043; IHC: fold-change 2.6 ± 0.9, P = 0.014). The proportion of proliferating GCG^+^ cells detected by flow cytometry and immunohistochemistry at 2.8 mM glucose was similar in 4/4 donors (Fig. 7e-h) as was the response to harmine in 3/4 donors (Fig. 7e & f) (FC: fold-change 22.4 ± 9.9, NS; IHC: foldchange 11 ± 4.2, NS), glucose in 3/4 donors (Fig. 7e & g) (FC: fold-change 1.5 ± 1, NS; IHC: fold-change 1.5 ± 0.4, P = 0.018) and HB-EGF in 4/4 donors (Fig. 7e & h) (FC: fold-change 1.3 ± 0.6, P = 0.032; IHC: fold-change 1 ± 0.6, NS). The proportion of proliferating CK19^+^ cells detected at 2.8 mM glucose was comparable between both techniques in 4/4 donors (Fig. 7i-l). However, a significant increase in proliferating CK19^+^ cells was detected by flow cytometry in 3/3 donors in response to harmine that was absent when assessed by immunohistochemistry (Fig. 7i & j) (FC: fold-change 18.5 ± 6.4, NS; IHC: fold-change 2.7 ± 1.3, NS). The response to 16.7 mM glucose was comparable in 4/4 donors (Fig. 7i & k) (FC: fold-change 1.4 ± 0.6, NS; IHC: fold-change 1.4 ± 0.6, NS) whereas the response to HB-EGF was similar in only 2/4 donors (Fig. 7i & l) (FC: fold-change 1.8 ± 0.4, P = 0.022; IHC: fold-change 5.4 ± 1.4, P = 0.01). Overall, these data reveal that although the percentage of proliferating GCG^+^ cells detected by flow cytometry and immunohistochemistry was comparable in all conditions tested, significant differences in the percentage of proliferating CPEP^+^ and CK19^+^ cells were observed.

## DISCUSSION

In this study we evaluated the capacity of harmine, glucose and HB-EGF to stimulate β-cell proliferation in isolated human islets using a multi-faceted approach involving intact and dispersed islet cultures, flow cytometry and immunohisto/cytochemistry and several proliferation and cell type-specific markers. Our data indicate that in the majority of islet batches human β cells respond to harmine and HB-EGF irrespective of the experimental approach, whereas the response to glucose is dependent on the donor and the assay used to assess proliferation. The relative abundance of proliferating non-β cells in human islet batches prompted us to assess the effects of these mitogens on α cells, CPEP^-^/GCG^-^ cells and CK19^+^ cells. We found that harmine potently increased the proliferation of α cells and CPEP^-^/GCG^-^ cells whereas HB-EGF mainly stimulated CPEP^-^/GCG^-^ cell proliferation including CK19^+^ cells.

Although the percentage of proliferating β cells detected by immunohistochemistry for Nkx6.1 or CPEP were significantly correlated in all treatment conditions (Supplementary Fig. S5 online), the fraction of EdU^+^ cells was markedly different. In general, the fraction of EdU^+^/Nkx6.1^+^ cells was lower than the corresponding EdU^+^/CPEP^+^ cells. This is reminiscent of a study by Aamodt et al. ^31^ in dispersed human islets, who scored only β cells that were both insulin^+^ and pancreatic and duodenal homeobox-1 (Pdx-1)^+^ and described a considerably lower level of β-cell proliferation in response to harmine and glucose compared to studies where all insulin^+^ (or CPEP^+^) cells were scored ^17–19,21,24–27,32^. This phenomenon may be explained in part by the fact that actively proliferating β-cells express lower levels of differentiation markers ^33^, which would decrease the sensitivity of detection of double labelling. Although CPEP is associated with functional β-cell differentiation, higher expression levels of CPEP compared to Nkx6.1 may allow for the perdurance of CPEP protein in proliferating β cells.

By performing EdU and Ki67 staining of intact islets by immunohistochemistry we found that the percentage of CPEP^+^ cells stained for each of these markers was comparable and significantly correlated in all treatment conditions (Supplementary Fig. S5 online). This result was surprising given that EdU was added to the culture medium throughout the 72-h exposure to mitogens. The expectation was that EdU should accumulate in more cells than express Ki67 at the end of the assay. The similar levels may be due to both EdU and Ki67 capturing all phases of the cell cycle. Taken together with the general absence of apoptosis in EdU^+^ cells, our data confirm that EdU^+^ cells are proliferating.

The DYRK1A inhibitor harmine increased human β-cell proliferation irrespective of the assay used, and also increased proliferation of GCG^+^, CPEP^-^/GCG^-^, and CK19^+^ cells in human islet batches. Whether the potency of harmine differs between cell types, however, is not known as the mitogenic effects of harmine were only assessed at 10 μM. Nevertheless, these data are consistent with previous studies demonstrating a potent mitogenic effect of harmine on human β-cell replication both in vitro and in vivo ^21,26^. Furthermore, the increase in proliferation of non-β cells in response to harmine is expected given the ubiquitous nature of DYRK1A signaling, and is consistent with several studies. Indeed, Wang et al. ^21^ showed that combined inhibition of DYRK1A and TGFβ signaling stimulates proliferation of CK19^+^ duct cells and endocrine a, δ and PP cells. Accordingly, the DYRK1A inhibitor 5-iodotubercidin stimulates both β- and α-cell proliferation ^19^. Finally, using single-cell mass cytometry Wang et al. ^20^ shown α cells to be considerably more responsive than β cells to the mitogenic effects of harmine. Further studies will be required to identify drugs that selectively target the β-cell for proliferation to advance β-cell mass accretion as a viable therapeutic approach to treat diabetes.

We found that the capacity of glucose to increase human β-cell proliferation was dependent on the donor and assay used. Glucose was inactive in the majority of donors in dispersed islets, whereas in intact islets β-cell proliferation was increased in most donors when assessed by immunohistochemistry but not by flow cytometry. In dispersed islets the loss of the cellular microenvironment and/or extended culture times (72 h) might contribute to decreased β-cell survival and proliferation ^28^. Our findings contrast with previous reports showing that glucose (15-20 mM) can increase β-cell proliferation to up to 2 % (approximately 2-fold over basal) in dispersed human islets cultured from 16-96 hours ^17,18,24,27,32^, but are in line with our previous immunohistochemical data in intact human islets ^34^. Although the underlying reasons for the differences in the β-cell proliferative response to glucose between our different assays ex vivo are unclear, it is noteworthy that variability in the response to glucose has been observed in vivo. Transplantation of human islets into streptozotocin-induced diabetic non-obese diabetic-severe combined immunodeficient mice revealed a correlation between glucose levels and β-cell proliferation following glucose infusion ^16^. Similarly, transplantation of human islets into spontaneously diabetic immunodeficient non-obese diabetic-Rag1null IL2rγnull Ins2Akita mice led to β-cell proliferation that positively correlated with glucose levels ^35^. Yet, when adult human islets were transplanted into NSG-DT mice, a model of hyperglycemia, no increase in β-cell proliferation was detected ^36^. In light of our ex vivo data, we speculate that the effect of glucose on human β-cell proliferation in vivo may be mostly indirect, whereas the direct effects of glucose, in striking contrast to the mouse β cell ^9,10,18,37^, may be relatively minor.

We recently demonstrated that the EGF receptor ligand HB-EGF exerts mitogenic activity on human β cells ^22^. In the present study we substantiated these findings by showing that β cells proliferate in response to HB-EGF in the majority of human islet batches in both intact and dispersed islets. Although α cells were only weakly responsive to HB-EGF, CPEP^-^/GCG^-^ cells and CK19^+^ ductal cells were highly responsive to HB-EGF. However, it is not known whether α cells respond to the mitogenic effects of HB-EGF at different concentrations since HB-EGF was only tested at 100 ng/ml. In agreement with these findings, the EGF receptor is expressed in β, α and ductal cells in the adult human pancreas ^38,39^. Furthermore, ductal cells constitute a neogenic source of β cells in rodents and humans ^40^ that are also responsive to EGF receptor ligands. In mice, adenoviral transduction of HB-EGF leads to ductal cell proliferation and differentiation into insulin^+^ cells ^41^. In humans, ductal cells proliferate in response to EGF and the related ligand betacellulin ^39^, and the combination of EGF and gastrin promotes ductal cell expansion and differentiation into Pdx-1^+^/insulin^+^ cells ^42^. Hence, the proliferating β cells detected in the presence of HB-EGF in this study could arise from replication of existing β cells and/or β-cell neogenesis from ductal cells.

Discrepancies between the various approaches used in the present study and with previous reports point to the sensitivity of human β cells to the culture conditions, for example using dispersed or intact islets, but also the need for rigorous labeling and scoring of proliferating β cells. In particular, the abundance of non-β-cell types in human islet batches could lead to the mis-identification of some proliferating cells as β cells. Importantly, we note that non-β cell types are rarely scored in human islet cell proliferation assays yet human islet batches contain multiple cell types including other endocrine cells but also ductal and acinar as well as endothelial, immune and stellate cells ^29^.

By comparing immunohistochemistry and flow cytometry to assess proliferation we found that the percentage of proliferating GCG^+^ cells was highly correlated in all conditions tested, whereas no significant correlation was detected in the percentage of proliferating CPEP^+^ and CK19^+^ cells (Supplementary Fig. S5 online). We identified a number of advantages and disadvantages of immunostaining and flow cytometry that may account for these differences. In our experience accurately determining the cell type identity of a nucleus in images of dispersed islet cells and intact islet sections is difficult, even when the operator is blinded to the experimental conditions, which limits the reliability of immunostaining approach. In contrast, flow cytometry eliminates the subjectivity of visual scoring and a higher number of cells can be easily and much more rapidly counted, increasing confidence in the data. However, the need for more extensive manipulations in the processing of samples for flow cytometry increases the level of cell death/loss compared to immunostaining. We conclude that multiple markers for β cells (e.g. INS, CPEP, Nkx6.1, Pdx-1) and proliferating cells (e.g. EdU, Ki67, phospho-histone H3) as well as endocrine, exocrine and other cell types should be stained simultaneously or in parallel to enhance the robustness of the conclusions and assess the relative efficacy of potential β-cell mitogens on several islet cell populations. We hope that these findings will provide useful guidance to reliably assess human islet cell proliferation ex vivo in the quest to identify novel and selective β-cell mitogens for potential therapeutic applications.

## METHODS

### Human islets

Islets from non-diabetic human donors were provided by the Alberta Diabetes Institute Islet Core, the Clinical Islet Laboratory at the University of Alberta, and the National Institute of Diabetes and Digestive and Kidney Diseases-funded Integrated Islet Distribution Program (IIDP; Supplementary Table S1 online). The use of human islets was approved by the Institutional Ethics Committee of the Centre Hospitalier de l’Université de Montréal (protocol # ND-05-035). All experiments were performed in accordance with the Institutional Ethics Committee guidelines and regulations. Informed consent was obtained from all donors and/or their legal guardians. Human islets were cultured in cGMP Prodo Islet Media (Standard) with 5 % (vol./vol.) human AB serum and 1 % glutamine/glutathione mixture (Prodo Labs Inc., Irvine, CA) overnight prior to experimentation.

### Measurement of cell proliferation in dispersed islets by immunocytochemistry

Dispersed human islet proliferation was assessed essentially as described ^22^. Briefly, hand-picked human islets were dispersed in accutase for 10 min at 37°C and plated in 96-well plates (Perkin Elmer Inc., Waltham, MA) treated with Poly-D-Lysine Hydrobromide (Sigma-Aldrich, St. Louis, MO). The next day, cells were cultured in RPMI-1640 (Thermo Fisher Scientific, Waltham, MA) with 1 % (vol./vol.) human serum albumin (Celprogen, Torrance, CA) for 3 days in the presence of EdU (10 μM) and glucose (2.8, 5.5 or 16.7 mM), harmine (10 μM; Millipore-Sigma, Darmstadt, Germany) or HB-EGF (100 ng/ml; R&D Systems, Minneapolis, MN) as indicated in the Figure legends. At the end of the treatment, cells were fixed with 3.7 % paraformaldehyde and immunostained for Nkx6.1 (Supplementary Table S2 online). The proliferative marker EdU was detected using the Click-iT EdU Imaging Kit with Alexa Fluor 594 (Thermo Fisher Scientific). Images were acquired with an Operetta CLS high-content imaging system (Perkin Elmer Inc., Waltham, MA) at 20X magnification using Harmony 4.5 software (https://www.perkinelmer.com/en-ca/product/operetta-cls-system-hh16000000). Cell counts were determined using ImageJ software (National Institutes of Health; https://imagej.net/Fiji). Proliferation was calculated as the percentage of double-positive EdU^+^/Nkx6.1^+^ cells over the total Nkx6.1^+^ population. Between 1,000 and 1,500 Nkx6.1+ cells from three different wells were manually counted per condition for each experimental replicate, as indicated in the Figure legends.

### Measurement of cell proliferation in intact islets by immunohistochemistry

Hand-picked human islets (approximately 200/condition) were cultured in RPMI-1640 with 1 % (vol./vol.) human serum albumin in 6-well plates for 3 days in the presence of glucose (2.8 or 16.7 mM), HB-EGF (100 ng/ml) or harmine (10 μM) as indicated in the Figure legends. EdU (10 μM) was added throughout. Media were replaced daily. At the end of treatment, islets were embedded in optimal cutting temperature (OCT) compound, frozen, sectioned at 8 μm and mounted on SuperFrost Plus slides (Thermo Fisher Scientific). Sections were fixed with 3.7 % paraformaldehyde and immunostained for Nkx6.1, CPEP, GCG or CK19 (Supplementary Table S2 online). Proliferation was detected by immunostaining for Ki67 (Supplementary Table S2 online) or EdU using the Click-iT EdU Imaging Kit with Alexa Fluor 594 or Pacific Blue.

Secondary antibodies were from Jackson ImmunoResearch (West Grove, PA). Images were taken with an Axio Imager upright fluorescence microscope with ZEN 2012 imaging software (Zeiss, Oberkochen, Germany; https://www.zeiss.com/microscopy/int/products/microscope-software/zen.html). Between 1,000 and 1,500 Nkx6.1^+^, CPEP^+^, GCG^+^ or CK19^+^ cells from 7-21 islets on 2 separate slides were manually counted per condition for each experimental replicate, as indicated in the Figure legends.

### Measurement of cell proliferation in intact islets by flow cytometry

At the end of the treatments described above, islets were dispersed in Accutase (1 μl/islet; Innovative Cell Technologies, Inc., San Diego, CA) for 10 min at 37°C and dead cells were labeled using the LIVE/DEAD Fixable Aqua (405 nm) Dead Cell Stain Kit (BD Biosciences, San Jose, CA). EdU detection, using the Click-iT Plus EdU Flow Cytometry Assay Kit with Alexa Fluor 488, and immunostaining were performed according to the manufacturer’s instructions (Thermo Fisher Scientific, Waltham, MA). Fluorophore-coupled primary antibodies and dilutions are listed in Supplementary Table S2 online. Flow cytometry analysis was carried out using a LSRIIB flow cytometer with BD FACSDiva software (BD Biosciences, San Jose, CA). Data was analyzed using FlowJo v10.7 software (Ashland, OR; https://www.flowjo.com/solutions/flowjo). Dead-cell stain, EdU-, CPEP-, GCG-, and CK19-labelled cells were detected using the 405-, 488-, 640- and 561-nm lasers coupled with 525/50-, 530/30-, 670/14- and 586/15-nm BP filters, respectively. Proliferation was calculated as the percentage of double-positive cells for EdU and each population marker over the corresponding total cell population for that marker.

### Measurement of apoptosis in intact islets

Apoptotic cells were visualized with the terminal deoxynucleotidyl transferase (TdT)-FragEL DNA Fragmentation Detection Kit (Millipore, Etobicoke, ON, Canada) according to the manufacturer’s instructions.

### Statistics

Significance was tested using two-tailed paired t-tests or ordinary one-way ANOVA, using GraphPad Prism 9 software (San Diego, CA; https://www.graphpad.com/scientific-software/prism/). P < 0.05 was considered significant. Fold-change data are expressed as mean ± SEM.

## DATA AVAILABILITY

All data generated or analyzed during this study are included in this published article (and its Supplementary Information files).

## ACKNOWLEDGMENTS

This study was supported by the National Institutes of Health (grant R01-DK-58096 to V.P.), the Canadian Institutes of Health Research (grant MOP 77686 to V.P.), and the Cardiometabolic Health, Diabetes and Obesity research Network of Quebec. Human pancreatic islets were provided by the IIDP (RRID:SCR _014387) at City of Hope, NIH Grant # 2UC4DK098085 and the JDRF-funded IIDP Islet Award Initiative. H.M. was supported by a doctoral studentship from the Fonds de Recherche Québec - Santé. V.P. holds the Canada Research Chair in Diabetes and Pancreatic B Cell Function.

## AUTHOR CONTRIBUTIONS

J.G., H.M. and V.P. designed the experiments, analyzed the results, and wrote the manuscript. H.M., J.G., and C.T. acquired the data. All authors revised the manuscript and approved the final version. V.P. is the guarantor of this work and, as such, takes full responsibility for the work.

## ADDITIONAL INFORMATION

The authors have no relevant conflict of interest to disclose.

