## Supplementary Tables and Figures for "Pronounced proliferation of non-beta cells in response to beta-cell mitogens in isolated human islets of Langerhans"

**Supplementary Figure S1. Analysis of apoptosis in intact human islets.** Intact human islets (N = 1) were exposed to 2.8 mM glucose (Vehicle or 2.8 Glu), harmine (10 μM), 16.7 mM glucose or HB-EGF (100 ng/ml) for 72 h. EdU (10 μM) was added throughout. Proliferation was assessed by EdU staining and apoptosis using the terminal deoxynucleotidyl transferase (TdT) FragEL DNA fragmentation assay. Representative images of FragEL (green), EdU (red) and nuclei (blue) staining for each condition are shown. Arrows highlight rare $EdU^+$/$FragEL^+$ nuclei. Scale bar, 50 μm. Images were acquired using an Axio Imager with ZEN 2012 software (https://www.zeiss.com/microscopy/int/products/microscope-software/zen.html).

**Supplementary Figure S2. Analysis of total and proliferating C-peptide- and Nkx6.1-positive cells in intact human islets.** Intact human islets (N = 3) were exposed to 2.8 mM glucose (Vehicle or 2.8 Glu), harmine (10 μM, Har), 16.7 mM glucose (16.7 Glu) and HB-EGF (100 ng/ml, HB) for 72 h. EdU (10 μM) was added throughout. C-peptide (CPEP) and Nkx6.1 were used to mark β cells and EdU for proliferation. Representative images of CPEP (green), Nkx6.1 (red) and EdU (blue) staining are shown (a). Arrows and arrowheads highlight $EdU^+$/$CPEP^+$/$Nkx6.1^-$ and $EdU^+$/ $CPEP^+$/$Nkx6.1^+$ cells, respectively. Scale bar, 50 μm. Images were acquired using an Axio Imager with ZEN 2012 software (https://www.zeiss.com/microscopy/int/products/microscope-software/zen.html). A minimum of 1,500 $CPEP^+$ cells were counted for each sample. The ratio of total (b) and proliferating (c) Nkx6.1$^+$/CPEP$^+$ cells for each condition are shown. Graphs were generated using GraphPad Prism 9 software (https://www.graphpad.com/scientific-software/prism/).

**Supplementary Figure S3. Analysis of CK19-positive cells in intact human islets.** Intact human islets (N = 2) were stained for C-peptide (CPEP, green), cytokeratin-19 (CK19, red) and nuclei (blue). Images of islet sections from 2 donors are shown. Scale bar, 50 µm. Images were acquired using an Axio Imager with ZEN 2012 software (https://www.zeiss.com/microscopy/int/products/microscope-software/zen.html).

**Supplementary Figure S4. Analysis of cell proliferation in intact human islets by flow cytometry.** Intact human islets were exposed to 2.8 mM glucose, harmine (10 µM), 16.7 mM glucose and HB-EGF (100 ng/ml) for 72 h. EdU (10 µM) was added throughout. At the end of the treatment islets were dispersed into single cells and analyzed by flow cytometry for C-peptide (CPEP), glucagon (GCG) and EdU (a-c & e-g) or cytokeratin 19 (CK19) and EdU (d & h). Representative plots showing gating used for selection of islet cells (a), live cells (a), CPEP$^+$, GCG$^+$, CPEP$^-$/GCG$^-$ (c) and CK19$^+$ (d) cells and proliferating EdU$^+$/GCG$^+$ (e), EdU$^+$/CPEP$^+$ (f), EdU$^+$/CPEP$^-$/GCG$^-$ (g) and EdU$^+$/CK19$^+$ (h) cells. Plots were generated using FlowJo v10.7 software (https://www.flowjo.com/solutions/flowjo).

**Supplementary Figure S5. Comparison of the percentage of proliferating cells detected using different approaches.** (a-e) Intact human islets were exposed to 2.8 or 16.7 mM glucose, harmine (10 µM) or HB-EGF (100 ng/ml) for 72 h. EdU (10 µM) was added throughout. Proliferation was assessed in parallel for each donor by immunohistochemistry (IHC) (a-e) or flow cytometry (FC) (c-e) and presented as a percentage of EdU$^+$/CPEP$^+$ over total CPEP$^+$ (a-c), EdU$^+$/Nkx6.1$^+$ over total Nkx6.1$^+$ (a), Ki67$^+$/CPEP$^+$ over total CPEP$^+$ (b), EdU$^+$/GCG$^+$ over total GCG$^+$ (d) and EdU$^+$/CK19$^+$ over total CK19$^+$ (e). Scatterplots of all treatment conditions combined are presented to illustrate the correlation between β-cell proliferation assessed by IHC for CPEP versus Nkx6.1 (6 donors) (a) and Ki67 versus EdU (3 donors) (b) and proliferation of CPEP$^+$ (5 donors) (c), GCG$^+$ (4 donors) (d) and CK19$^+$ (4 donors) (e) cells assessed by IHC versus FC. These data are presented in the main figures 2 and 3 (a), 3 and 4 (b) and 7 (c-e). Graphs were generated using GraphPad Prism 9 software (https://www.graphpad.com/scientific-software/prism/). Significance was tested using two-tailed paired t-tests. $P<0.05$ was considered significant. NS, not significant. R, Pearson correlation coefficient.

Supplementary table S1.xlsx

Supplementary table S2.docx

**Supplementary Figure S1**

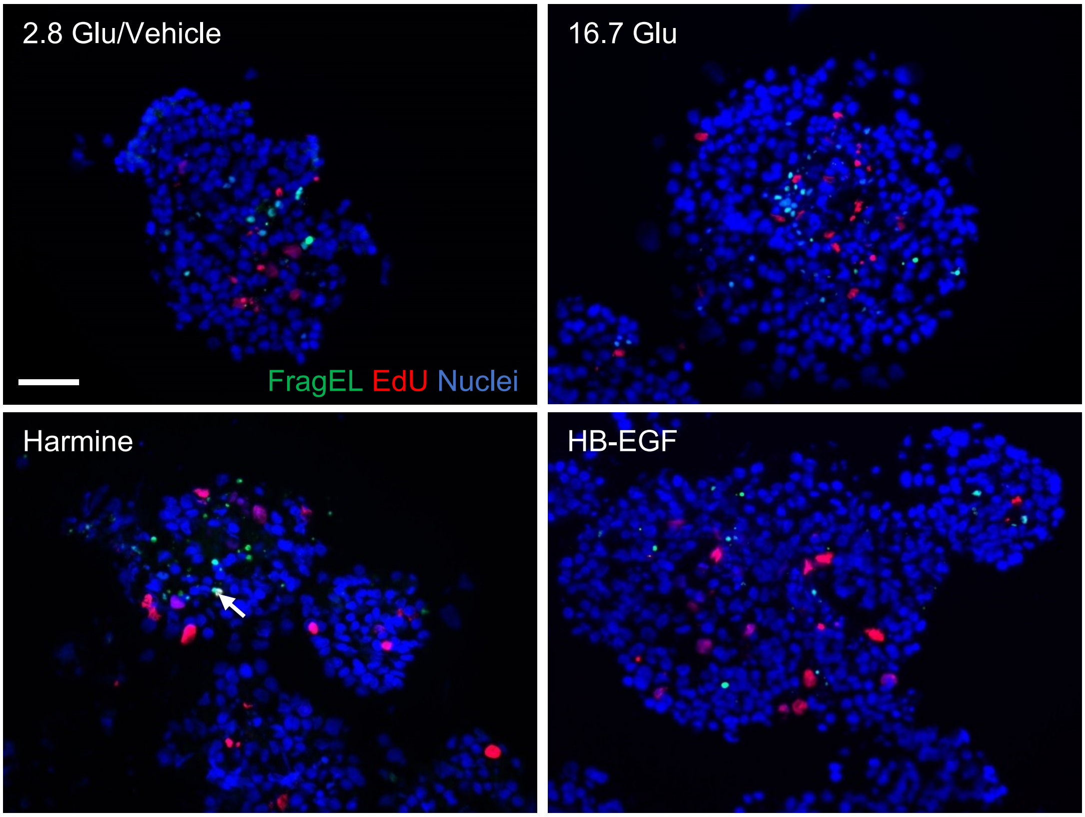

#### Supplementary Figure S2

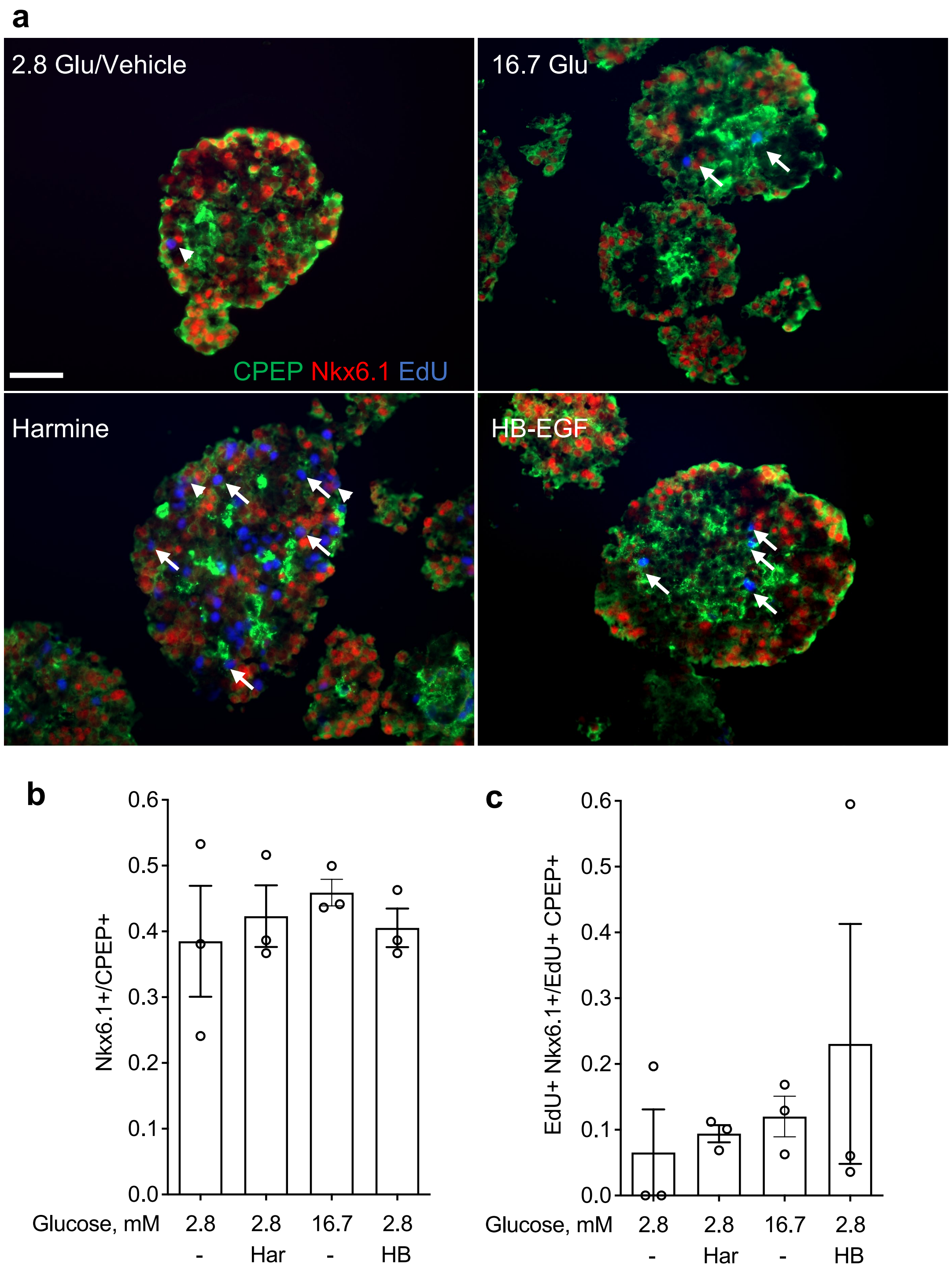

### Supplementary Figure S3

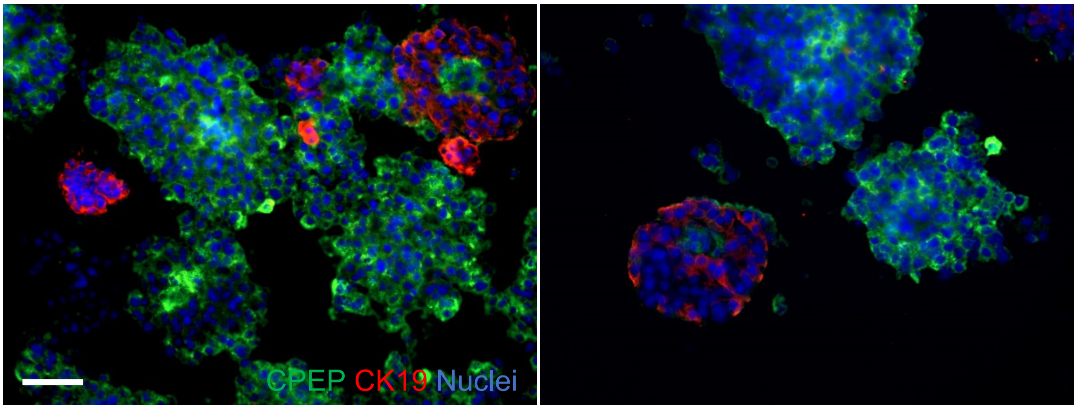

### Supplementary Figure S4

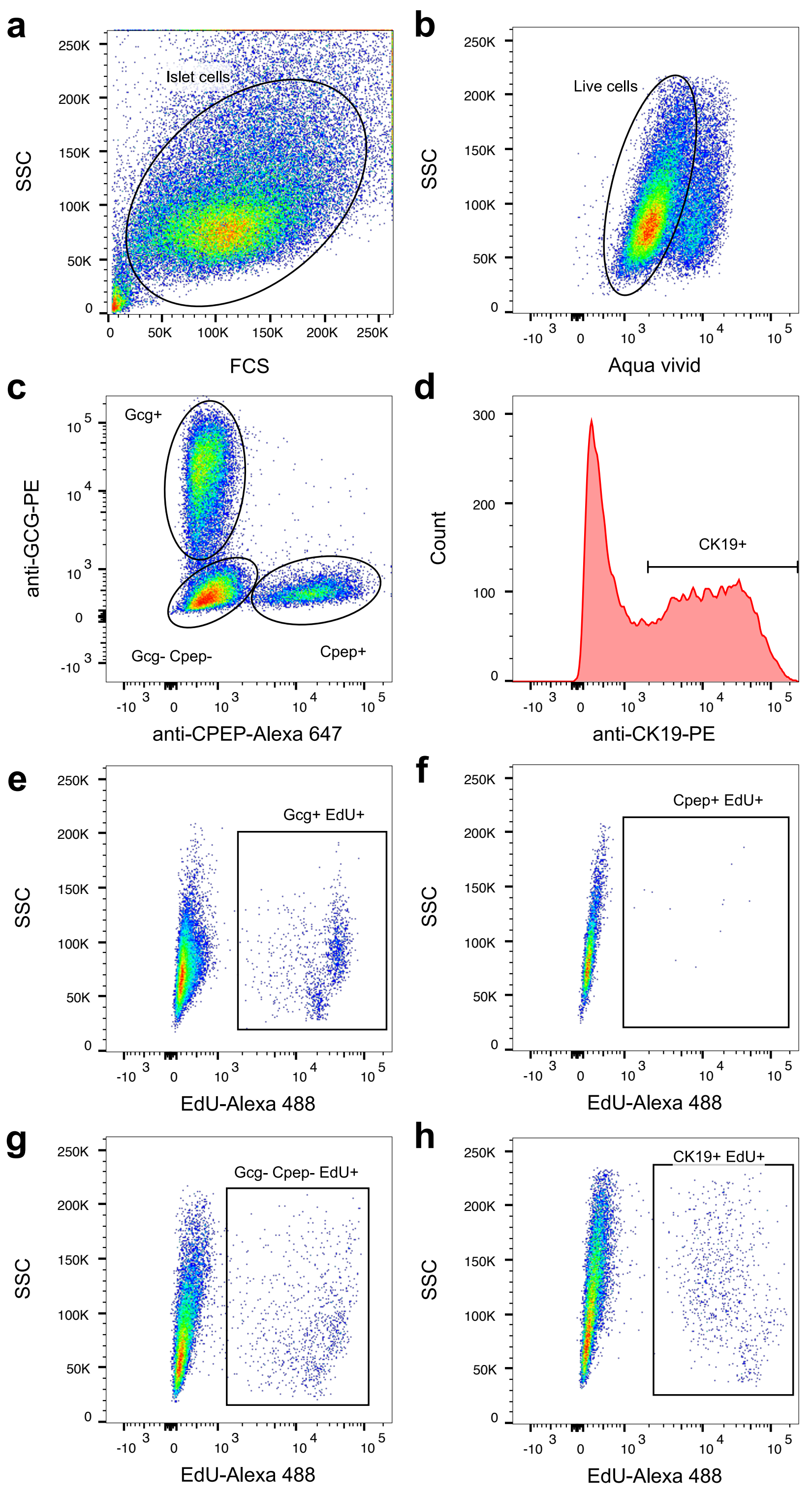

#### Supplementary Figure S5

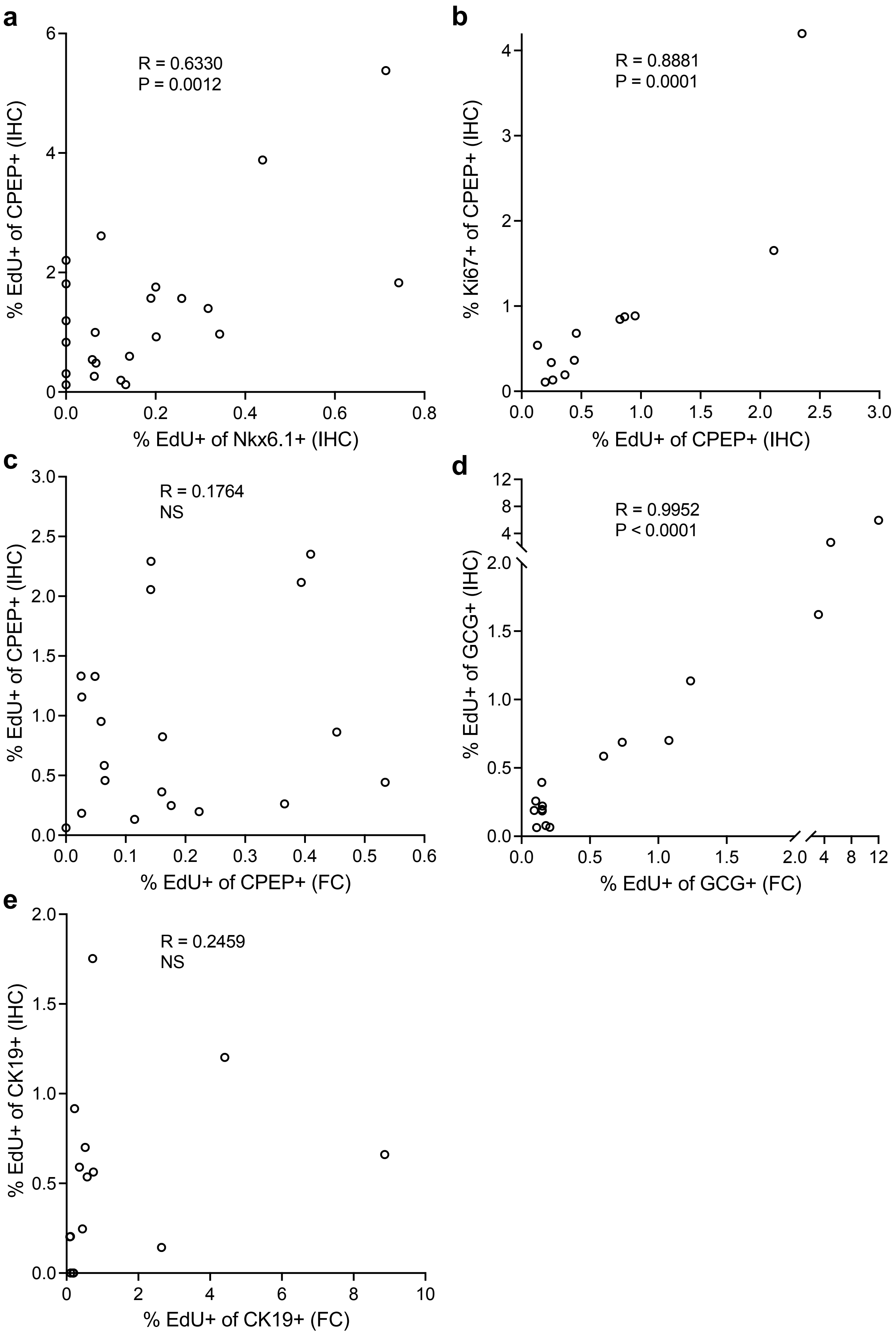

| Supplementary Table S1. Human islet donors used in study. | | | | | | | | | |
|---|---|---|---|---|---|---|---|---|---|
| **Donor** | **1** | **2** | **3** | **4** | **5** | **6** | **7** | **8** | **9** |
| **Unique identifier** | R340 | SAMN12924398 | SAMN13027809 | R346 | SAMN13134368 | R348 | R341 | R344 | SAMN11791244 |
| **Donor Age (years)** | 36 | 38 | 43 | 27 | 62 | 43 | 42 | 59 | 47 |
| **Donor Sex (M/F)** | M | M | F | M | M | F | M | M | M |
| **Donor BMI (kg/m$^2$)** | 23.3 | 28.0 | 31.6 | 24.1 | 31.3 | 16.4 | 30 | 27.8 | 36.1 |
| **Donor HbA1c** | 5.3 | 5.4 | 5.7 | 5.7 | 5.5 | 5.7 | Not Reported | 5.5 | 5.7 |
| **Origin/source of islets** | Alberta IsletCore | IIDP | IIDP | Alberta IsletCore | IIDP | Alberta IsletCore | Alberta IsletCore | Alberta IsletCore | IIDP |
| **Islet isolation center** | Alberta IsletCore | The Scharp-Lacy Research Institute | The Scharp-Lacy Research Institute | Alberta IsletCore | The Scharp-Lacy Research Institute | Alberta IsletCore | Alberta IsletCore | Alberta IsletCore | Southern California Islet Cell Resource Center |
| **Donor history of diabetes** | No | No | No | No | No | No | No | No | No |
| **Cause of death** | NDD-Neurological | Head trauma | Anoxia | NDD-Neurological | Cerebrovascular/stroke | NDD-Neurological | NDD-Neurological | NDD-Neurological | Cerebrovascular/stroke |
| **Warm ischemia time (h)** | Not Reported | Not Reported | 0.2 | Not Reported | Not Reported | Not Reported | Not Reported | Not Reported | 0.3 |
| **Cold ischemia time (h)** | 12.5 | 8.8 | 11.1 | 12 | 10.2 | 15.5 | 15.75 | 11.5 | 6.4 |
| **Estimated purity (%)** | 95 | 80 | 90 | 90 | 90 | 90 | 95 | 95 | 95 |
| **Estimated viability (%)** | | 95 | 95 | | 95 | | | | 96 |
| **Glucose-stimulated insulin secretion stimulation index (SI)** | SI (G2.8mM-G16.7mM)= 4.56 | SI (G2.8mM-G28mM)= 1.8 | SI (G2.8mM-G28mM)= 2.8 | SI (G2.8mM-G16.7mM)= 6.17 | Not Reported | SI (G2.8mM-G16.7mM)= 1.63 | SI (G2.8mM-G16.7mM)= 9.81 | SI (G2.8mM-G16.7mM)= 3.11 | SI (G2.8mM-G28mM)= 2.5 |
| **FACS data** | | | | | | | | | |
| **Single cells/total cells (%)** | | | | | | | | | |
| **Live cells/single cells (%)** | | | | | | | | | |
| **CPEP+ cells/live cells (%)** | | | | | | | | | |
| **GCG+ cells/live cells (%)** | | | | | | | | | |
| **CPEP-, GCG- cells/live cells (%)** | | | | | | | | | |

| | | | | | | | | | |
|---|---|---|---|---|---|---|---|---|---|
| **<u>Donor</u>** | **10** | **11** | **12** | **13** | **14** | **15** | **16** | **17** | **18** |
| **Unique identifier** | R301 | SAMN10869303 | SAMN10023853 | SAMN09862214 | SAMN09370567 | SAMN10410585 | H2182 | SAMN09929594 | SAMN10516338 |
| **Donor Age (years)** | 18.0 | 25.0 | 25.0 | 31.0 | 32.0 | 38.0 | 48.0 | 51.0 | 52.0 |
| **Donor Sex (M/F)** | M | M | F | M | M | M | F | F | F |
| **Donor BMI ($kg/m^2$)** | 19 | 27.7 | 33.5 | 31.8 | 28.5 | 28.2 | 38.5 | 29.7 | 26.9 |
| **Donor HbA1c** | 5.0 | 5.2 | 5.1 | 5.3 | 5.2 | 5.9 | | 5.8 | 4.5 |
| **Origin/source of islets** | Alberta IsletCore | IIDP | IIDP | IIDP | IIDP | IIDP | Alberta islet lab | IIDP | IIDP |
| **Islet isolation center** | Alberta IsletCore | The Scharp-Lacy Research Institute | Southern California Islet Cell Resource Center | Southern California Islet Cell Resource Center | University of Wisconsin | The Scharp-Lacy Research Institute | Alberta islet lab | Southern California Islet Cell Resource Center | The Scharp-Lacy Research Institute |
| **Donor history of diabetes** | No | No | No | No | No | No | No | No | No |
| **Cause of death** | NDD-Neurological | Anoxia | Anoxia | Head trauma | Head trauma | Head trauma | | Cerebrovascular/strok e | Cerebrovascular/strok e |
| **Warm ischemia time (h)** | Not Reported | 0.5 | 0.1 | Not Reported | 0.1 | Not Reported | Not Reported | Not Reported | Not Reported |
| **Cold ischemia time (h)** | 5 | 8.4 | 7.0 | 5.5 | 11.5 | 7.3 | | 6.9 | 6.3 |
| **Estimated purity (%)** | 75 | 90 | 75 | 80 | 95 | 90 | | 75 | 80 |
| **Estimated viability (%)** | | 95 | 96 | 95 | 91 | 95 | | 94 | 95 |
| **Glucose-stimulated insulin secretion stimulation index (SI)** | SI (G2.8mM-G16.7mM)= 9.21 | SI (G2.8mM-G28mM)= 8.9 | SI (G2.8mM-G28mM)= 0.7 | SI (G2.8mM-G28mM)= 2.0 | SI (G2.8mM-G28mM)= 1.8 | SI (G2.8mM-G28mM)= 6.6 | Not Reported | SI (G2.8mM-G28mM)= 2.1 | SI (G2.8mM-G28mM)= 6.5 |
| **<u>FACS data</u>** | | | | | | | | | |
| **Single cells/total cells (%)** | 87.6 | 89.6 | 85.0 | 93.9 | 95.6 | 93.2 | 89.6 | 96.1 | 92.5 |
| **Live cells/single cells (%)** | 88.9 | 86.5 | 85.2 | 83.2 | 88.5 | 74.7 | 77.9 | 91.1 | 83.5 |
| **CPEP+ cells/live cells (%)** | 8.6 | 43.2 | 23.4 | 3.1 | 6.7 | 19.3 | 19.3 | 9.2 | 13.7 |
| **GCG+ cells/live cells (%)** | 6.9 | 35.8 | 18.3 | 28.9 | 62.1 | 29.9 | 12.9 | 55.6 | 44.2 |
| **CPEP-, GCG- cells/live cells (%)** | 84.5 | 21.0 | 58.3 | 68.0 | 31.2 | 50.8 | 67.8 | 35.2 | 42.1 |

| | | | | | | | | | |
|---|---|---|---|---|---|---|---|---|---|
| **Donor** | **19** | **20** | **21** | **22** | **23** | **24** | **25** | **26** | **27** |
| **Unique identifier** | R305 | R271 | SAMN09228907 | R356 | HP20071-01 | SAMN15337453 | R365 | H2330 | SAMN16081314 |
| **Donor Age (years)** | 60.0 | 60.0 | 62 | 45 | 42 | 26 | 31 | 49 | 24 |
| **Donor Sex (M/F)** | M | F | M | F | M | M | F | F | M |
| **Donor BMI (kg/m$^2$)** | 21 | 25.9 | 36.1 | 29.7 | 27.97 | 24.2 | 20.3 | 27.2 | 20.1 |
| **Donor HbA1c** | 5.6 | 5.5 | 5.8 | 5.1 | 4.9 | | 4.8 | | 5.4 |
| **Origin/source of islets** | Alberta IsletCore | Alberta IsletCore | IIDP | Alberta IsletCore | Prodo labs | IIDP | Alberta IsletCore | University of Alberta | IIDP |
| **Islet isolation center** | Alberta IsletCore | Alberta IsletCore | The Scharp-Lacy Research Institute | Alberta IsletCore | Prodo labs | The Scharp-Lacy Research Institute | Alberta IsletCore | Alberta islet distribution program | The Scharp-Lacy Research Institute |
| **Donor history of diabetes** | No | No | No | No | No | No | No | No | No |
| **Cause of death** | NDD-Neurological | NDD-Neurological | Anoxia | NDD-Neurological | Head trauma | Head trauma | NDD-Neurological | Not Reported | Head trauma |
| **Warm ischemia time (h)** | Not Reported | Not Reported | Not Reported | Not Reported | Not Reported | Not Reported | Not Reported | Not Reported | Not Reported |
| **Cold ischemia time (h)** | 10 | 13.5 | 10.7 | 14 | Not Reported | 10.2 | 6.75 | | 7.8 |
| **Estimated purity (%)** | 80 | 95 | 90 | 80 | Not Reported | 98 | 95 | 72.5 | 95 |
| **Estimated viability (%)** | | | 95 | | | 95 | | 85.5 | 95 |
| **Glucose-stimulated insulin secretion stimulation index (SI)** | SI (G2.8mM-G16.7mM)= 1.29 | SI (G2.8mM-G16.7mM)= 4.98 | SI (G2.8mM-G28mM)= 3.9 | SI (G2.8mM-G16.7mM)= 5.49 | Not Reported | SI (G2.8mM-G28mM)= 4.4 | SI (G2.8mM-G16.7mM)= 5.34 | Not Reported | Not Reported |
| **FACS data** | | | | | | | | | |
| **Single cells/total cells (%)** | 91.4 | 98.6 | 92.6 | | | | | | 88.4 |
| **Live cells/single cells (%)** | 85.5 | 91.7 | 87.6 | | | | | | 84.3 |
| **CPEP+ cells/live cells (%)** | 31.1 | 34.2 | 2.6 | | | | | | 17.8 |
| **GCG+ cells/live cells (%)** | 14.2 | 37.0 | 83.1 | | | | | | 56.8 |
| **CPEP-, GCG- cells/live cells (%)** | 54.8 | 28.8 | 14.4 | | | | | | 25.5 |

| | | | | | | | | |
|---|---|---|---|---|---|---|---|---|
| **Donor** | **28** | **29** | **30** | **31** | **32** | **33** | **34** | **35** |
| **Unique identifier** | SAMN16114998 | R391 | SAMN16871526 | SAMN17277513 | SAMN17528599 | SAMN17928660 | SAMN18021384 | SAMN18092805 |
| **Donor Age (years)** | 24 | 67 | 62 | 43 | 60 | 62 | 56 | 56 |
| **Donor Sex (M/F)** | F | M | M | F | M | M | M | M |
| **Donor BMI ($kg/m^2$)** | 45.5 | 24.5 | 21.0 | 36.5 | 29.9 | 25.9 | 24.2 | 21.6 |
| **Donor HbA1c** | 5.2 | 4.9 | 5.3 | 5.2 | 5.8 | 5.1 | 5.8 | 5.1 |
| **Origin/source of islets** | IIDP | Alberta IsletCore | IIDP | IIDP | IIDP | IIDP | IIDP | IIDP |
| **Islet isolation center** | University of Wisconsin | Alberta IsletCore | The Scharp-Lacy Research Institute | Southern California Islet Cell Resource Center | University of Pennsylvania | Southern California Islet Cell Resource Center | The Scharp-Lacy Research Institute | Southern California Islet Cell Resource Center |
| **Donor history of diabetes** | No | No | No | No | No | No | No | No |
| **Cause of death** | Anoxia | Not Reported | Cerebrovascular/stroke | Cerebrovascular/stroke | Anoxia | Head trauma | Anoxia | Cerebrovascular/stroke |
| **Warm ischemia time (h)** | Unknown | Not Reported | Not Reported | 0.4 | Not Reported | Not Reported | Unknown | Not Reported |
| **Cold ischemia time (h)** | 4.6 | 17 | 9.7 | 5.3 | 7.7 | 7.4 | 6.7 | 5.8 |
| **Estimated purity (%)** | 85 | 95 | 90 | 90 | 85 | 90 | 90 | 90 |
| **Estimated viability (%)** | 98 | | 95 | 96 | 98 | 96 | 95 | 96 |
| **Glucose-stimulated insulin secretion stimulation index (SI)** | Not Reported | SI (G2.8mM-G16.7mM)=2.31 | SI (G2.8mM-G28mM)= 1.1 | SI (G2.8mM-G28mM)= 2.7 | SI (G2.8mM-G28mM)= 1.3 | Not Reported | Not Reported | Not Reported |
| **FACS data** | | | | | | | | |
| **Single cells/total cells (%)** | 89.3 | 91.4 | 91.6 | 86.4 | 86.5 | 92.9 | | |
| **Live cells/single cells (%)** | 83.4 | 94.5 | 93.6 | 90.5 | 91.0 | 90.4 | | |
| **CPEP+ cells/live cells (%)** | 16.6 | 41.6 | 15.9 | 16.5 | 32.9 | 12.6 | | |
| **GCG+ cells/live cells (%)** | 54.7 | 31.3 | 72.2 | 46.8 | 49.3 | 67.7 | | |
| **CPEP-, GCG- cells/live cells (%)** | 28.7 | 27.1 | 11.8 | 36.6 | 17.8 | 19.6 | | |

**Supplementary Table S2** Source of antibodies used in study.

| Antibody | Dilution/Concentration | Company | Cat # |
|---|---|---|---|
| **Immunohistochemistry/ Immunocytochemistry** | | | |
| Rabbit polyclonal Anti-Nkx6.1 | 1:100 | Novus | NBP1-82553 |
| Rat monoclonal Anti-C-Peptide | 5 μg/ml | DSHB | GN-ID4 |
| Mouse monoclonal Anti-Cytokeratin 19 | 1:500 | Abcam | Ab9220 |
| Mouse monoclonal Anti-Glucagon | 1:400 | Abcam | Ab10988 |
| Rabbit polyclonal Anti-Ki67 | 1:400 | Abcam | Ab15580 |
| **Flow cytometry** | | | |
| Alexa Fluor® 647 Mouse Anti-C-Peptide | 1:50 | BD Biosciences | 565831 |
| PE Mouse Anti-Glucagon | 1:50 | BD Biosciences | 565860 |
| Mouse monoclonal Anti-Cytokeratin 19 | 1:500 | Abcam | Ab9220 |
